# Maternal obesity alters human milk oligosaccharides content and correlates with early acquisition of late colonizers in the neonatal gut microbiome

**DOI:** 10.1101/2025.05.09.653164

**Authors:** Karina Corona-Cervantes, Víctor H. Urrutia-Baca, July S. Gámez-Valdez, Brenda Jiménez-López, Nora A. Rodríguez-Gutierrez, Karla Chávez-Caraza, Francisca Espiricueta-Candelaria, Ulises A. Salas Villalobos, Perla Ramos-Parra, Janet A. Gutierrez Uribe, Marion Brunck, Cristina Chuck Hernández, Cuauhtemoc Licona-Cassani

## Abstract

The infant gastrointestinal tract is critical for metabolic and immune development, shaped by early gut microbiome colonization. Maternal obesity may disrupt this process by altering human milk oligosaccharides (HMOs), influencing microbial succession and infant health. This longitudinal study examines maternal BMI-associated variations in HMOs and infant fecal microbiota. Human milk samples from 97 mothers (stratified by BMI) were collected at 48h, one month, and three months postpartum, with HMOs quantified via UPLC-MS/MS. In a subsample of this population, infant fecal microbiome composition was analyzed using metagenomics tools. Mothers with obesity exhibited reduced 2’-FL, LNnT, 3’-SL, 6’-SL, LNFPI concentration at 48 h postpartum. Infants born to these mothers showed diminished early colonizers such as Gammaproteobacteria at 48h, alongside lower antigen and polyamine biosynthesis pathways. Conversely, late colonizers like *Lachnospiraceae* species were more abundant at one and three months postpartum. Sialylated and neutral HMOs negatively correlated with these late colonizers, which metabolize such oligosaccharides. These findings suggest maternal obesity alters HMO profiles, potentially disrupts pioneer microbial establishment and accelerating late-colonizer dominance. Reduced keystone bacteria and shifted metabolic pathways may impair immune priming, with long-term health implications. This highlights the interplay between maternal BMI, HMO composition, and infant microbiome succession, underscoring the need to address maternal metabolic health to optimize early microbial colonization and immune programming.

## Introduction

The early neonatal period is a critical window for metabolic and immune development, driven by dynamic interactions between the infant gut microbiome and host (Selma-Royo et al. 2020; Catassi et al. 2024; Kalbermatter et al. 2021). During the first week of life, the neonatal gut microbiome undergoes dynamic shifts, beginning with pioneer taxa like *Enterobacteria* and *Staphylococcus*, which thrive in the oxygen-rich neonatal gut nutrient availability (Milani et al. 2017). These earliest colonizers shape the microenvironment for late anaerobic successors such as *Bifidobacterium* and *Bacteroides* (Houghteling and Walker 2015). This early microbial succession is not merely structural but functionally plays a crucial role in the ecosystem development and immune system programming (Jian et al. 2021; Sprockett et al. 2018). Microbial colonization in infancy shapes immune tolerance, barrier integrity, and metabolic programming, with lifelong implications for disease susceptibility (Al Nabhani and Eberl 2020; Kalbermatter et al. 2021).

During the neonatal period, human milk oligosaccharides (HMOs) play a central role in shaping the microbiome. These complex neutral or acidic molecules, composed of lactose linked to fucosylated or sialylated monosaccharides, function as both prebiotics and immunomodulators (Atyeo and Alter 2021; Carr et al. 2021; Rousseaux et al. 2021; Singh et al. 2022). Structurally diverse HMOs selectively nourish keystone commensals such as *Bifidobacterium* and *Bacteroides*, which metabolize these glycans into bioactive molecules that modulate immune responses and support cross-feeding interactions among microbial species (Maessen et al. 2020). Additionally, HMOs contribute to gut immunity by enhancing barrier function and acting as glycan decoys to prevent pathogen colonization. By promoting the growth of certain microbes while inhibiting others, HMOs serve as key modulators of gut microbial ecology (Ojima et al. 2022)

Maternal factors, such as obesity, can alter human milk composition. For instance, obese mothers exhibit reduced HMO levels and lower Proteobacteria abundance in colostrum (Gámez-Valdez et al. 2021; Cortés-Macías et al. 2023; Saben et al. 2021; Urrutia-Baca et al. 2024) . Disruption of the colonization pattern may have long-term consequences, as the premature acquisition of *Clostridia* and *Lachnospiraceae*, has been linked to immune dysregulation and metabolic disorders (Sbihi et al. 2019; Shenhav et al. 2024) . Longitudinal data are essential to capture the time-dependent dynamics of microbial succession and to disentangle whether maternal obesity alters HMO profiles in a stage-specific manner during the first 3 months.

Here, we conducted a longitudinal study of 97 Mexican mother-infant dyads, profiling HMO concentrations, and gut microbiota composition and metagenomic functionality over in sub sample of infants in the first three months of life. We compared HMO profiles in normal weight (NW) *vs* mothers with overweight or obesity (OW/OB) and track bacterial succession dynamics in a sub sample of infants. This work addresses a gap in understanding how maternal obesity disrupts the milk-microbiome axis—a key driver of infant immune and metabolic programming. By integrating longitudinal HMO and metagenomic data, we provide insights into how early colonization defects may predispose infants to lifelong health risks, to create actionable strategies to mitigate intergenerational cycles of metabolic disease in vulnerable populations, particularly in regions with high obesity prevalence, such as Mexico.

## Results

### Demographic and clinical characteristics of the study cohort

A total of 97 volunteer women agreed to participate in the present study after meeting the eligibility criteria. The women were divided into two study groups, 50 women with normal weight and 47 women with overweight and obesity. Sociodemographic, anthropometric data, and mother-child clinical histories were obtained, and colostrum samples within the first 48 hours after delivery (M0) were collected from all of them and six meconium samples were collected from their babies. For longitudinal follow-up of the participants and their babies, a home visit was made in the first month postpartum (M1) where 17 women with normal weight and 25 women with overweight or obesity decided to continue in the present study; anthropometric data, mature milk sample at M1 and 12 stool samples from their babies were collected. At the third month after delivery (M3), a final home visit was made where 15 women with normal weight and 19 women with overweight or obesity decided to continue in the present study; anthropometric data, mature milk samples at M3 and 8 stool samples from their babies were collected (Figure 1).

**Figure 1.**
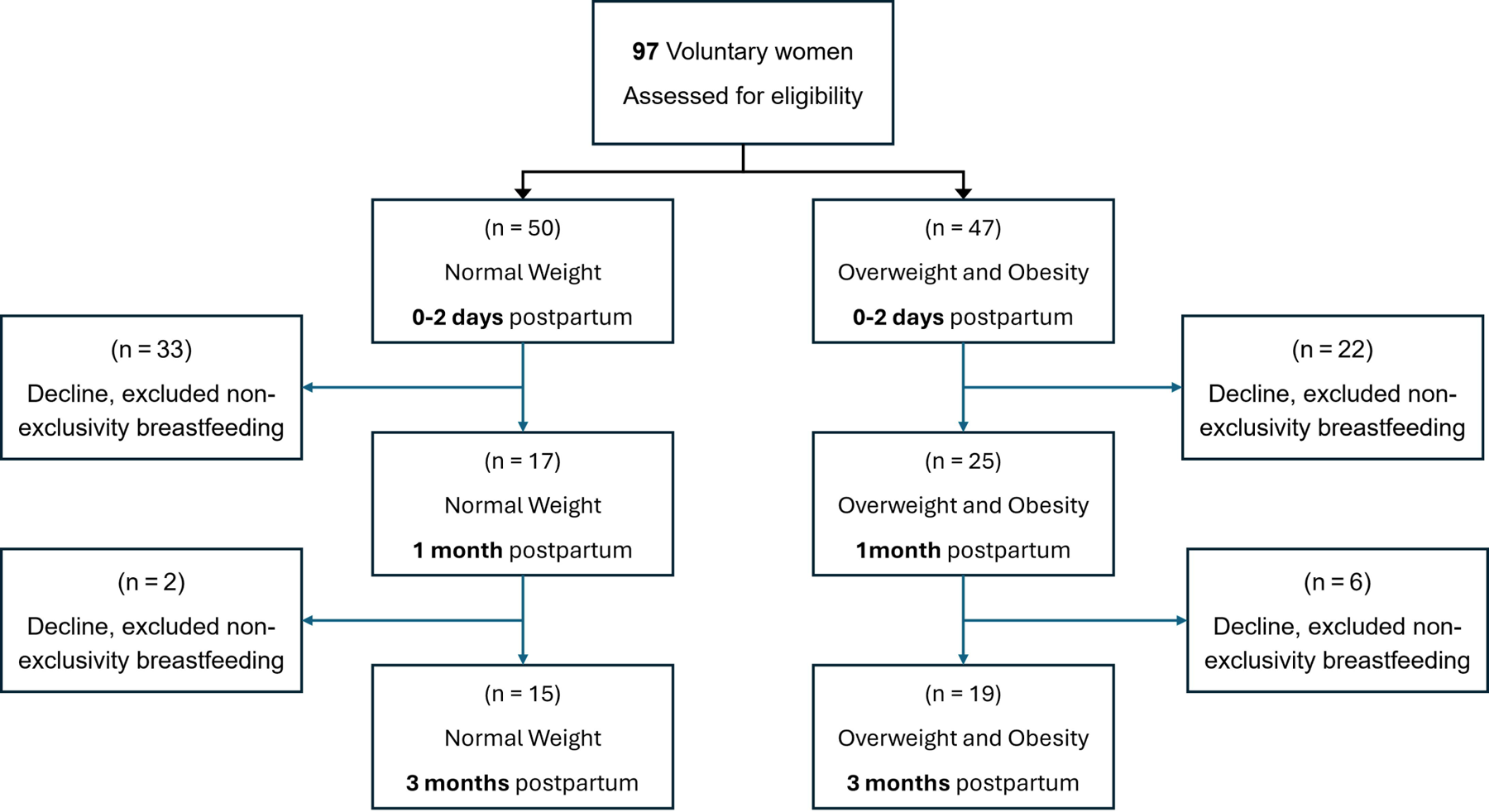
Flow diagram for participant recruitment, follow up, and analysis for HMOs in milk samples at the long of the study.

Participating mothers had an average age of 23.56 ± 5.74 years at delivery (Table S1). Regarding their demographics, 82.5% resided in the Monterrey N.L. Mexico metropolitan area, 63.9% had middle school as their highest level of completed education, and 87.6% of the mothers were employed in various unspecified occupations. Medically, 10.3% had a history of high blood pressure and 4.1% of cholesteremia or dyslipidemia; none had a history of kidney disease. Cholesteremia or dyslipidemia was found to be associated with overweight or obesity in mothers (p = 0.035); the four women who had this medical history were in the overweight or obesity group, as detailed in Table S1. Furthermore, the diastolic and systolic blood pressures in all participants were 69.7 ± 10.7 mm Hg and 113.9 ± 11.8 mm Hg, respectively. Systolic blood pressure was significantly higher (p = 0.005) in women with overweight or obesity (116.6 ± 9.1 mm Hg) compared to women with normal weight (111.5 ± 13.4 mm Hg). Based on the anthropometric data in all participants, the pre-pregnancy Body Mass Index (BMI) was 23.7 ± 8.9 kg m^-2^, and the BMI at delivery was 28.9 ± 6.4 kg m^-2^ with a total weight gain of 10.2 ± 6.5 kg. Regarding the data associated with pregnancy and childbirth, 68.0% of births were spontaneous at the beginning, and 69.1% of the deliveries were eutocic according to their conclusion; each delivery resulted in the birth of a single child. The average number of previous births was 2.1 ± 1.3.

Data were obtained from all participating mothers’ newborns: 57.7% were female, and 96.9% did not show congenital anomalies. The weight, height and BMI at birth were 3.2 ± 0.4 kg, 49.4 ± 2.1 cm and 12.9 ± 1.3 kg m^-2^, respectively. In addition, the average birth head circumference was 33.7 ± 1.4 cm. Regarding birth weight, height, birth head circumference, sex, and congenital anomalies, no differences were found between the groups (p > 0.05). About z-scores for infant growth assessment, length-for-age, weight-for-age, weight-for-length and head circumference-for-age were −0.2 ± 1.1, −0.3 ±0.9, −0.4 ± 1.1, −0.4 ± 1.3 and −1.2 ± 5.0, respectively. In the weight-for-age z-score, statistical differences (p =0.035) were observed between children born to women with overweight/obesity (−0.1±0.8) and those born to women with normal weight (−0.5±0.9); no differences were observed between the groups in the other z-score values. Regarding neonatal screening, APGAR and Silverman-Andersen scores were normal in most newborns, and hearing screening was positive (a positive result indicates correct hearing at birth) in 51.5% (Table S1). In longitudinal follow-up, no differences were observed between the groups in the anthropometric parameters of the mothers or their children at the first and third month postpartum, as shown in Table S1.

### Maternal Clinical and Biochemical Factors Are Linked to HMO Concentrations and Infant Growth

We analyzed correlations between maternal factors (BMI, age, parity, mode of delivery, blood pressure) and HMO concentrations (neutral core, fucosylated, sialylated) using a linear mixed model. Biochemical markers including glucose, triglycerides, creatinine, and uric acid were also assessed in relation to HMO levels in our cohort.

Our results show BMI is inversely correlated with concentrations of sialylated and neutral core HMOs, while maternal age showed negative associations specifically with neutral, non-fucosylated HMOs (Figure S1 and Table S2). Biochemical markers such as glucose, triglycerides, and total protein were positively linked to various HMO types, notably Lacto-N-neotetraose (LNnT), which exhibited broad associations across these parameters (Table S3). Infant growth, assessed through WHO metrics at one and three months postpartum, revealed positive associations between maternal glucose and indirect bilirubin levels with infant growth metrics, whereas urea was linked to weight-for-length and zBMI (Table S4). HMOs like Lacto-N-Fucopentaose I (LNFPI) and 3-Fucosyllactose (3-FL) positively influenced infant length metrics but showed a negative association with infant weight early on, while sialylated and fucosylated HMOs supported infant size and length by three months, highlighting their beneficial role in infant growth.

### Maternal BMI Significantly Reduces HMO Concentrations in Colostrum but not in Later Lactation Stages

This study focuses exclusively on secretor mothers. Analysis of seven HMOs across three lactation stages revealed that in human milk, fucosylated oligosaccharides, particularly 2’ Fucosyllactose (2’-FL) and 3-FL are the most abundant HMOs. These were followed by neutral HMOs, such as Lacto-N-Tetraose (LNT) and LNFPI, and sialylated HMOs, including 3’-Sialyllactose (3’-SL) and 6’-Sialyllactose (6’-SL), as shown in Figure 2A. We noted that some colostrum samples had higher LNT levels, while others were richer in 3’-SL. To further explore the variability in HMO concentrations, we grouped mothers by BMI, categorizing them into normal weight and obesity groups (Figure 2B).

**Figure 2.**
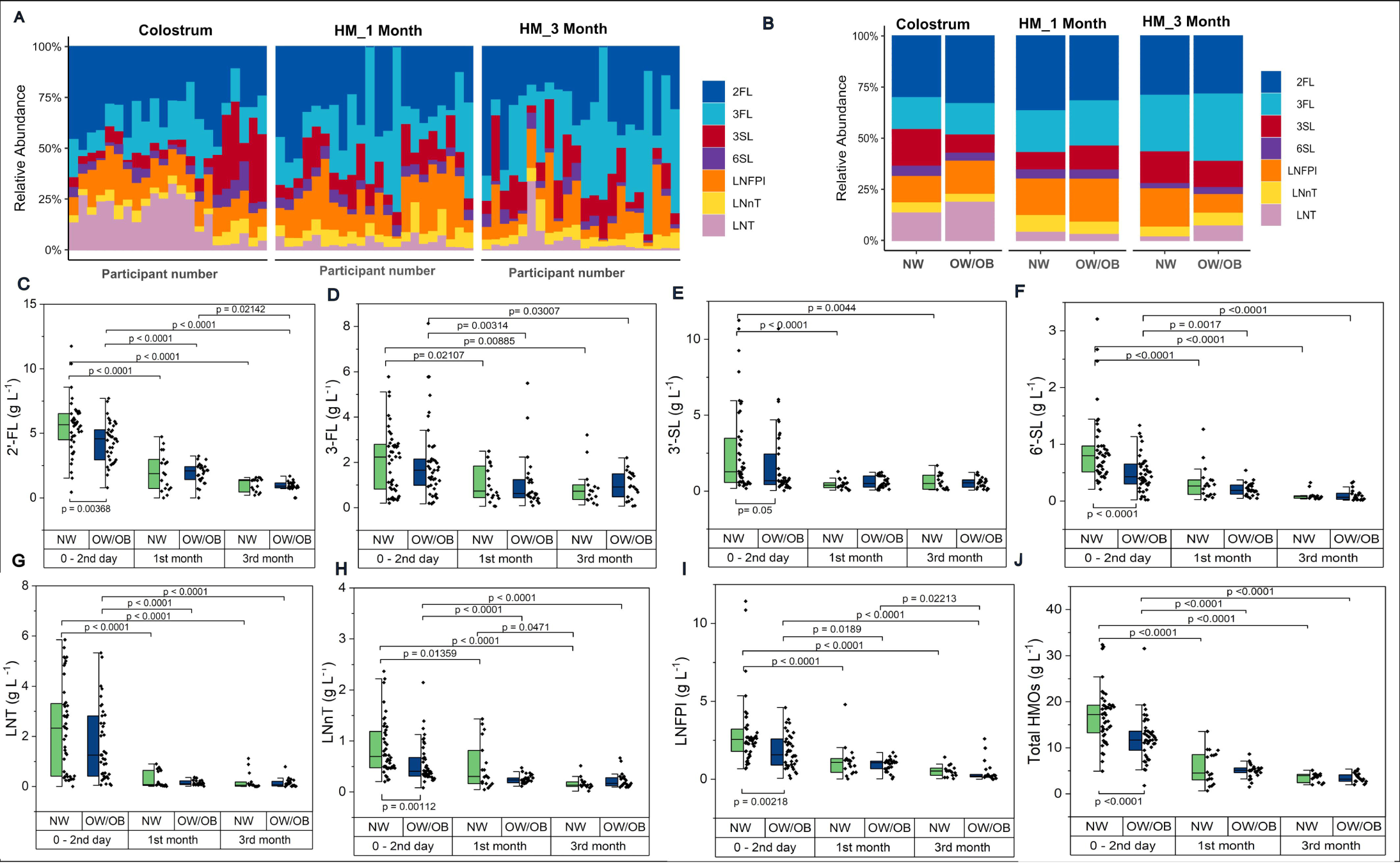
*HMOs concentrations of longitudinal milk samples at three points during lactation*. (A) Individual HMO abundance for each participant mother (B) HMO abundance according BMI groups at the 0-2 days, 1st and 3rd month postpartum. (C-J) Differences between the study from Mexican mothers with normal weight and overweight/obesity. (C) 2’-fucosyllactose, (D) 3-fucosyllactose, (E) lacto-N-fucopentaose I, (F) 3’-sialyllactose, (G) 6’-sialyllactose, (H) lacto-N-neotetraose, (I) lacto-N-tetraose and and (J) Total HMOs. The data is displayed in a grouped boxplot and only p values < 0.05 are shown. The p values of the top were obtained using the non-parametric Kruskal–Wallis test for the evaluation of the differences between the lactation period. The p values of the bottom were obtained using the non-parametric Mann–Whitney U test for the evaluation of the differences between the study groups. NW: women with normal weight, OW/OB: women with overweight/obesity.

The median concentration of total HMOs in the groups were significantly different (p <0.0001), 17.2 g L^-1^ (IQR= 13.1-19.3) and 11.7 g L^-1^ (IQR= 8.9-13.7) in the colostrum of women with normal weight and women with overweight or obesity, respectively. No differences were observed between groups in mature milk at M1 and M3. However, the median concentration of total HMOs decreased significantly (p <0.0001) in mature milk at M1 (4.5 g L^-1^, IQR=2.8-8.8) and M3 (3.9 g L^-1^, IQR=2.4-4.2) respect to colostrum at M0 in women with normal weight and women with overweight or obesity (M1: 5.1 g L^-1^, IQR=4.5-5.6 and M3: 3.2 g L^-1^, IQR=2.7-4.2), as shown in (Figure 2C).

Significant differences were observed in the median concentrations of 2’-FL (NW: 5.7 g L^-1^, IQR=4.5-6.5 ***vs.*** OW/OB: 4.6 g L^-1^, IQR=2.9-5.4; p =0.00368), 3’-SL (NW: 1.27 g L^-1^, IQR=0.6-3.5 ***vs.*** OW/OB: 0.7 g L^-1^, IQR=0.4-2.6; p =0.0465), 6’-SL (NW: 0.8 g L^-1^, IQR=0.5-0.9 ***vs.*** OW/OB: 0.4 g L^-1^, IQR=0.2-0.7; p <0.0001), LNFPI (NW: 2.6 g L^-1^, IQR=1.8-3.2 ***vs.*** OW/OB: 1.6 g L^-1^, IQR=0.9-2.6; p =0.00218) and LNnT (NW: 0.7 g L^-1^, IQR=0.5-1.2 ***vs.*** OW/OB: 0.4 g L^-1^, IQR=0.3-0.7; p =0.00112) between the groups but only at M0. The median concentration in 6 of the 7 oligosaccharides studied decreased significantly at M1and M3 postpartum respect to colostrum (M0) in both women with normal weight and women with overweight or obesity (Figure 2C-2J).

### Infant Gut Microbiome Composition Exhibits Temporal Shifts and Varies with Maternal Pre-Gestational BMI

To examinate how bacterial communities shift across developmental stages of the infant and to determine the impact of maternal pregestational BMI on microbiome composition, fecal samples were collected at three key time points at M0, M1 and M3

The taxonomic composition showed changes across different stages of early life. At 48 hours postpartum, the bacterial microbiome was dominated by Proteobacteria in comparison with M1 and M3 (*p* = 0.010), while at one month and three months post-partum, Actinobacteria become more prevalent (p= 0.0415) (Figure S2A). At the genus level, the M0 stage is characterized by a predominance of *Enterobacter* and *Klebsiella*, whereas M1 and M3 are marked by a higher presence of *Bifidobacterium* (Figure S2B).

When infants were grouped by maternal BMI, differences were observed at the phylum level. *Proteobacteria* were more abundant in the normal-weight group across all stages (M0, M1, M3) compared to infants born to mothers with obesity. In contrast, *Firmicutes* and *Actinobacteria* were significantly more abundant in the obesity group. These observed trends were not significantly different (Table S5).

Beta diversity analysis showed no significant differences between the microbiota of infants in the normal weight and obesity groups at any of the sampling points (Figure S3C-3E). These findings suggest the bacterial composition of the infant gut changes according to the maternal BMI particularly in the first hours after birth.

### Infant Gut Microbiome Diversity is Negatively Correlated with Sialylated and Neutral HMO Concentrations Across Developmental Stages

We evaluated the effect of maternal HMOs supply on infant gut microbiome diversity by assessing correlations between maternal oligosaccharide concentrations and both taxonomic and functional alpha diversity for each mother-infant pair. Taxonomic diversity was estimated using operational taxonomic units (OTUs), while functional diversity was determined based on the abundance of metabolic pathways in each sample.

Among 48 hours postpartum, higher concentrations of 3’-SL, LNT, and LNnT were associated with lower bacterial taxonomic richness, as indicated by the Chao1, observed species, and ACE indices (Figure 3A). Regarding functional pathway diversity, 6’-SL and LNnT were negatively correlated with the ACE richness index (Figure 3B). Notably, at this early neonatal stage, both taxonomic and functional diversity (Simpson, Shannon, and Fisher indices) were positively correlated with maternal pregestational BMI (Figure 3B).

**Figure 3.**
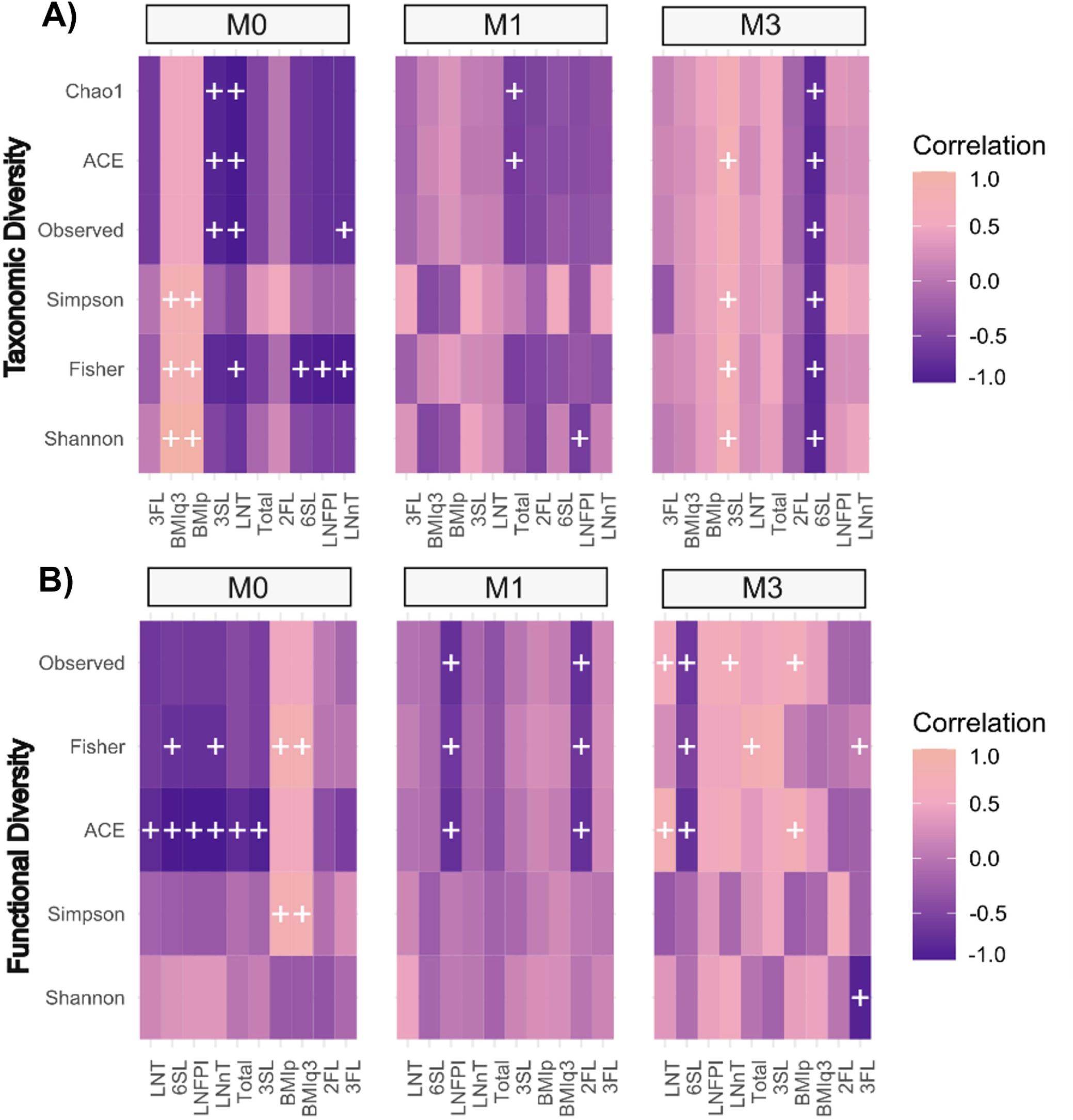
Longitudinal Associations Between Infant Gut Microbiome Diversity, Human Milk Oligosaccharide (HMO) Concentrations, and Maternal BMI at 0, 1, and 3 Months Postpartum. (A) Taxonomic and (B) functional alpha diversity indices of the infant gut microbiome in relation to maternal HMO concentrations and pregestational BMI. Associations were assessed using Spearman’s rank correlation analysis. Correlation heatmaps use a pink-to-dark purple gradient to indicate positive and negative associations, respectively. White (+) denotes p < 0.05, while gray (+) denotes p < 0.1.

By the first month, LNFPI was the only HMOs showing negative correlations with taxonomic Shannon diversity and functional richness (observed species, Fisher, and ACE indices) (Figures 3A and 3B). At three months postpartum, 6’-SL was significantly negatively correlated with all diversity indices. In contrast, 3’-SL showed positive correlations with bacterial richness (ACE index) and all diversity indices. Moreover, functional richness (based on observed species) was positively correlated with neutral core HMOs (LNT, LNnT) and maternal pregestational BMI at this stage (Figures 3A and 3B). These results highlight significant correlations between HMOs and the infant gut microbial community, reinforcing the role of HMOs in shaping gut microbiota by influencing bacterial richness and diversity across early developmental stages.

### Intermediate and late colonizers are increased in Infant Born from Mothers with Obesity are Associated with HMOs across diverse Growth stages

After establishing the relationship between infant gut microbiome diversity and maternal human milk oligosaccharides we investigated associations between specific differentially abundant based on maternal BMI at each developmental stage. To achieve this, we first identified the taxa that differed between infants born to mothers with normal weight and those born to mothers with obesity.

Our analysis revealed that most differences occurred at the M0 stage. Within 48 hours postpartum, neonates of mothers with obesity showed higher abundances of intermediate and late colonizers from the phyla Firmicutes, Actinobacteria, and Bacteroidetes. These included genera such as *Ligilactobacillus*, *Lactiplantibacillus*, *Lentilactobacillus*, *Eggerthella*, *Bifidobacterium*, *Bacteroides*, and *Akkermansia*. In contrast, neonates of normal-weight mothers had higher levels of bacteria from the phylum Proteobacteria, such as *Enterobacter*, *Klebsiella*, *Succinivibrio*, and *Escherichia* (Figure 4A).

**Figure 4.**
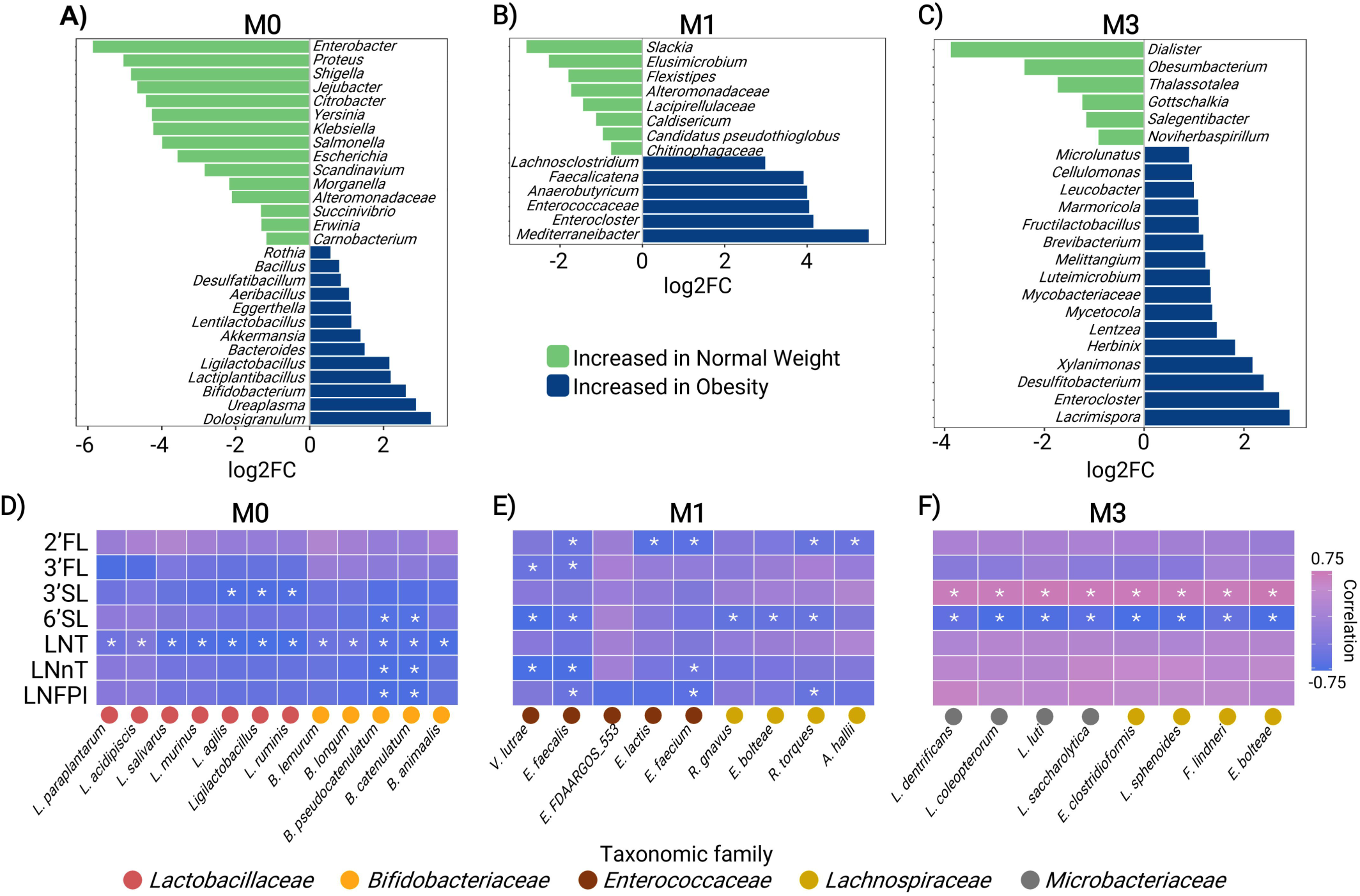
Associations Between Obesity-Related Differential Fecal Bacteria and Human Milk Oligosaccharide Concentrations Across Infant Growth Stages. (A–C) Differentially abundant bacterial genera in infants born to normal-weight mothers versus those born to mothers with obesity at three time points: (A) 0–2 days (M0), (B) 1 month (M1), and (C) 3 months (M3) postpartum. (D–F) Spearman’s rank correlation heatmaps showing the associations between differentially abundant genera and maternal HMO concentrations at (D) 0–2 days, (E) 1 month, and (F) 3 months postpartum. The left Y-axis represents HMO concentrations, while the right Y-axis represents the relative abundance of bacterial species.

After identifying the differential bacteria, we examined their associations with oligosaccharide concentrations in maternal milk thought Spearman correlation. Negative correlations between *Bifidobacterium* species and LNT, LNFPI, LNnT, and the sialylated HMO 6’-SL were observed. For instance, *B. pseudocatenulatum* and *B. longum* with all these oligosaccharides, *B. animalis* and *B. lemurum* for LNT, and *B. catenulatum* for 6’-SL (Figure 4D). Likewise, *Lactiplantibacillus* and *Ligilactobacillus* were negatively correlated with LNT and LNFPI. Specifically, *Ligilactobacillus ruminis* and *Ligilactobacillus agilis* were negatively associated with both LNT and 3-FL, while *Ligilactobacillus salivarius* and *Lactiplantibacillus plantarum* were correlated only with LNT (Figures 4D). In contrast, some genera that were more abundant in the normal weight group, such as *Enterobacter* and *Scandinavium*, were negatively correlated with 2’-FL.

Next, we evaluated the influence of maternal BMI on microbiome composition after one month postpartum. At this growth stage, an enrichment of late colonizers from the phylum Firmicutes in the infants of mothers with obesity. Particularly members of the family Lachnospiraceae, including the genera: *Mediterraneibacter*, *Enterocloster*, *Anaerobutyricum*, *Faecalitalea*, *Lacrimispora*, and *Lachnoclostridium*, as well as members of the family Enterococcaceae (Figure 4B). In contrast, genera such as *Slackia*, *Elusimicrobium*, *Flexispira*, and bacteria from the family *Chitinophagaceae* were more abundant in infants of mothers with normal weight.

Similarly to the M0 stage, species from genera enriched in infants born to mothers with obesity showed negative correlations with maternal oligosaccharides, particularly non-fucosylated oligosaccharides such as LNnT and 6’-SL. *Enterocloster boltae*, *Ruminococcus gnavus*, *Anaerobutyricum hallii*, *Vagococcus lutrae*, and *Enterococcus faecalis*, displayed negative correlations with 6’-SL, while *Ruminococcus torques*, *Enterococcus lactis*, and *Enterococcus faecalis* were negatively associated with 2’-FL. Meanwhile other species from the *Enterococcaceae* family, including *Enterococcus faecalis*, *E. faecium*, *FDAARGOS 553*, and *Vagococcus lutrae*, were negatively correlated with non-fucosylated neutral oligosaccharides, such as LNFPI and LNnT (Figures 4E). On the other hand, *Slackia heliotrinireducen* which was prevalent in infants of normal weight mothers, was positively correlated with 3’-SL and LNFPI. These findings suggest that maternal BMI and HMO composition continue to play a significant role in shaping the microbial landscape at this stage of growth.

The final growth stage evaluated was the third month postpartum. Here, similarly to the M1 stage, the microbiomes of infants from mothers with obesity were enriched with strict anaerobic bacteria members of the *Lacnospiraceae* family such as *Herbinix*, *Lacrimispora*, and *Enterocloster*, along with other genera as *Leucobacter*, *Cellulomonas*, *Brevibacterium*, *Microlunatus*, and *Fructilactobacillus.* Regarding the infants of normal-weight mothers*, Dialister*, *Obesumbacterium*, *Gottschalkia*, *Salegentilibacter*, and *Noviherbaspirillum* were more abundant (Figure 4C).

Here, similarly to the previous stages, bacteria associated with obesity showed negative correlations with 6’-SL. However, unlike earlier stages, these same bacteria display positive correlations with 3’-SL (Figure 4F). Specifically, species such as *Enterocloster bolteae*, *E. clostridioformis*, *Lacrimispora sphenoides*, *Lacrimispora saccharolytica*, and *Herbinix luporum* from the family Lachnospiraceae, along with *Leucobacter colepterorum*, *L. denitrificans*, *L. luti*, *Microlunatus elymi*, *Cellulomonas tanus*, *Brevibacterium linens*, and *Fructilactobacillus lindneri*, exhibited this pattern. Meanwhile, the genus *Gottschalkia*, which was more prevalent in infants of normal-weight mothers, also showed positive correlations with 3’-SL (Figure 4F).

These findings show that maternal oligosaccharides as sialylated HMOs are particularly correlated with taxa overrepresented in obesity at the long of the first three months post-partum, potentially impacting long-term microbial development.

### Polyamine Metabolic Pathways are Reduced in the Microbiome of Newborns from Mothers with Obesity Within the First 48 Hours Postpartum

The next step involved analyzing the infant metagenome, identifying a total of 4,200 genes mapped to 507 general metabolic pathways after processing the unstratified data. These pathways were categorized into superclasses based on the MetaCyc database: 234 related to biosynthesis, 141 to degradation, and 132 to precursor metabolism and energy production (encompassing processes such as fermentation, reduction, transformation, and shunting). We evaluated the relative abundance and variability of these pathways across all the fecal samples Subsequently, the relative abundances of the metabolic pathways were compared between groups based on maternal pregestational BMI at each stage of infant growth, revealing differences in the infants’ gut metagenome. Most of these differences were found in the microbiome of neonates at the first 48 hours postpartum stage, where 89 pathways were significantly different (p < 0.05, q < 0.05) between the groups. Of these, 68 pathways were more abundant in the normal weight group, while 21 were more prevalent in the obesity group. Complete lists of metabolic pathways associated with the gut metagenomes are available on Table S6.

In the normal-weight group, pathways involved in biosynthesis and metabolism of polyamine were significantly more abundant. These pathways primarily contribute to the production of putrescine and spermidine through the degradation of L-amino acids such as L-ornithine and L-arginine, as well as the biosynthesis of cadaverine via L-lysine degradation by decarboxylase enzymes (AST-PWY, POLYAMSYN-PWY, ARG+POLYAMINE-SYN, ORNDEG-PWY and PWY-6305) (Figure 5A Table S6). The enrichment of these metabolic pathways was primarily driven by the presence of *Enterobacter cloacae*, which was highly abundant in the neonatal microbiome of the normal-weight group and negatively correlated with both pregestational and post gestational maternal BMI (Figure 5B).

**Figure 5.**
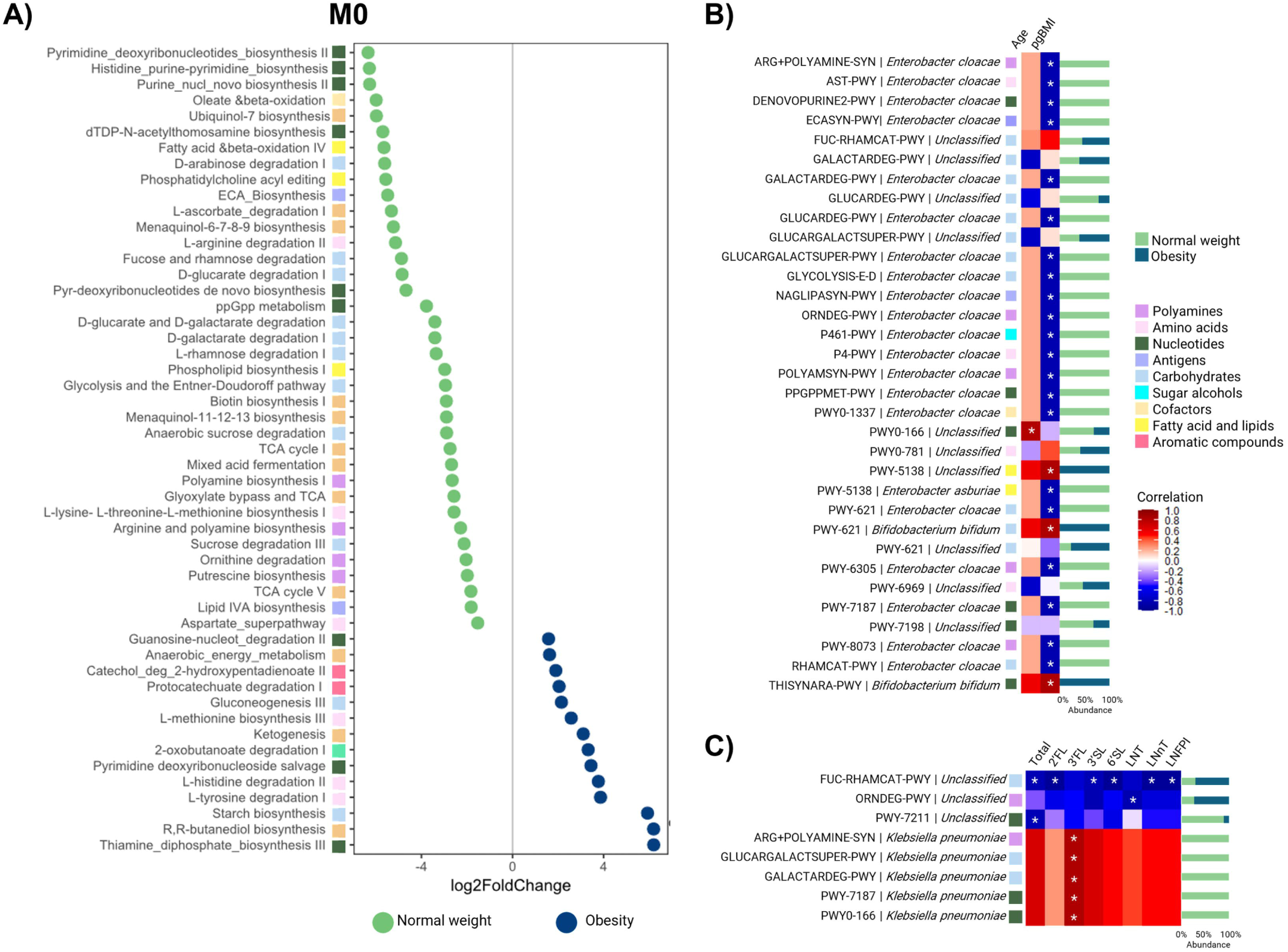
*Functional metabolic pathways at 48 hr postpartum in infant microbiome* A) Differential metagenomic pathways between infants born to normal weight and obese mothers. Correlation of microbiome metabolic pathways with B) Maternal variables and C) Human milk oligosaccharides concentration. The Spearman correlation index was employed to establish correlation. Blue shows negative correlation and red for positive.Stars represent a p value <0.05. Stacked Bar plots represent the relative abundance of the stratified pathway according to the maternal BMI.

In addition to polyamine biosynthesis, several other pathways were significantly enriched in the microbiome of infants born to normal-weight mothers and negatively correlated with maternal BMI. These included pathways involved in the biosynthesis of enterobacterial common antigens, such as lipopolysaccharides, phospholipids, and lipid IVA (ECASYN-PWY, PHOSLIPSYN-PWY, NAGLIPASYN-PWY, and PWY-8073) (Figure 5A), primarily represented in *Enterobacter cloacae* (figure 5B). Additionally, pathways related to carbohydrate degradation (e.g., D-galactarate, sucrose, D-glucarate, fucose, and L-rhamnose) and nucleic acid biosynthesis, particularly pyrimidine deoxyribonucleotides and purines, were notably prevalent in these neonates (Figure 5A-B).

In contrast, neonates born to mothers with obesity showed a greater representation of pathways related to the degradation of aromatic compounds, such as L-tyrosine and catechol. Additionally, their microbiome exhibited enrichment in carbohydrate metabolism, and generation of precursor metabolites and Energy pathways, including gluconeogenesis III, ketogenesis, and starch biosynthesis, as well as pathways involved in the biosynthesis of thiamine diphosphate, by *Bifidobacterium bifidum* (Figure 5A).

Next, we examined the association between stratified metabolic pathways and the concentrations of HMOs provided by the mother. We found that 3-FL was positively correlated with pathways abundant in infants of normal-weight mothers, including polyamine biosynthesis, D-galactarate degradation, and pyrimidine deoxyribonucleotide de novo biosynthesis, primarily by *Klebsiella pneumoniae* (Figure 5C). Conversely, pathways related to ornithine and fucose-rhamnose metabolism were negatively correlated with 2’-FL, sialylated HMOs, and non-fucosylated neutral HMOs, attributed to an unclassified bacterium (Figure 5B).

Given the pronounced enrichment of polyamine biosynthesis pathways in the metagenomes of neonates born to normal-weight mothers, we further investigated the differential abundance of genes encoding polyamine biosynthesis enzymes and transport systems. Our analysis revealed consistent enrichment of genes associated with polyamine metabolism and transport in the normal-weight group. Specifically, lysine decarboxylase—a key enzyme catalyzing cadaverine production from L-lysine—was significantly more abundant in this group (q < 0.05). Similarly, genes encoding enzymes for putrescine biosynthesis (via L-arginine and agmatine pathways) and spermidine synthase, which drives spermidine production, were elevated in the normal-weight cohort (Figure 6). Transport-related genes were also strongly represented in normal weight, including those encoding the arginine transport permease protein (K0999), antiporters mediating cadaverine-lysine (K03757) and putrescine-ornithine (K03756) exchange. Also, the spermidine/putrescine ABC transporter complex (*potA*, *potB*, *potC*, *potD*), and MdtI, a spermidine export protein (Figure 6).

**Figure 6.**
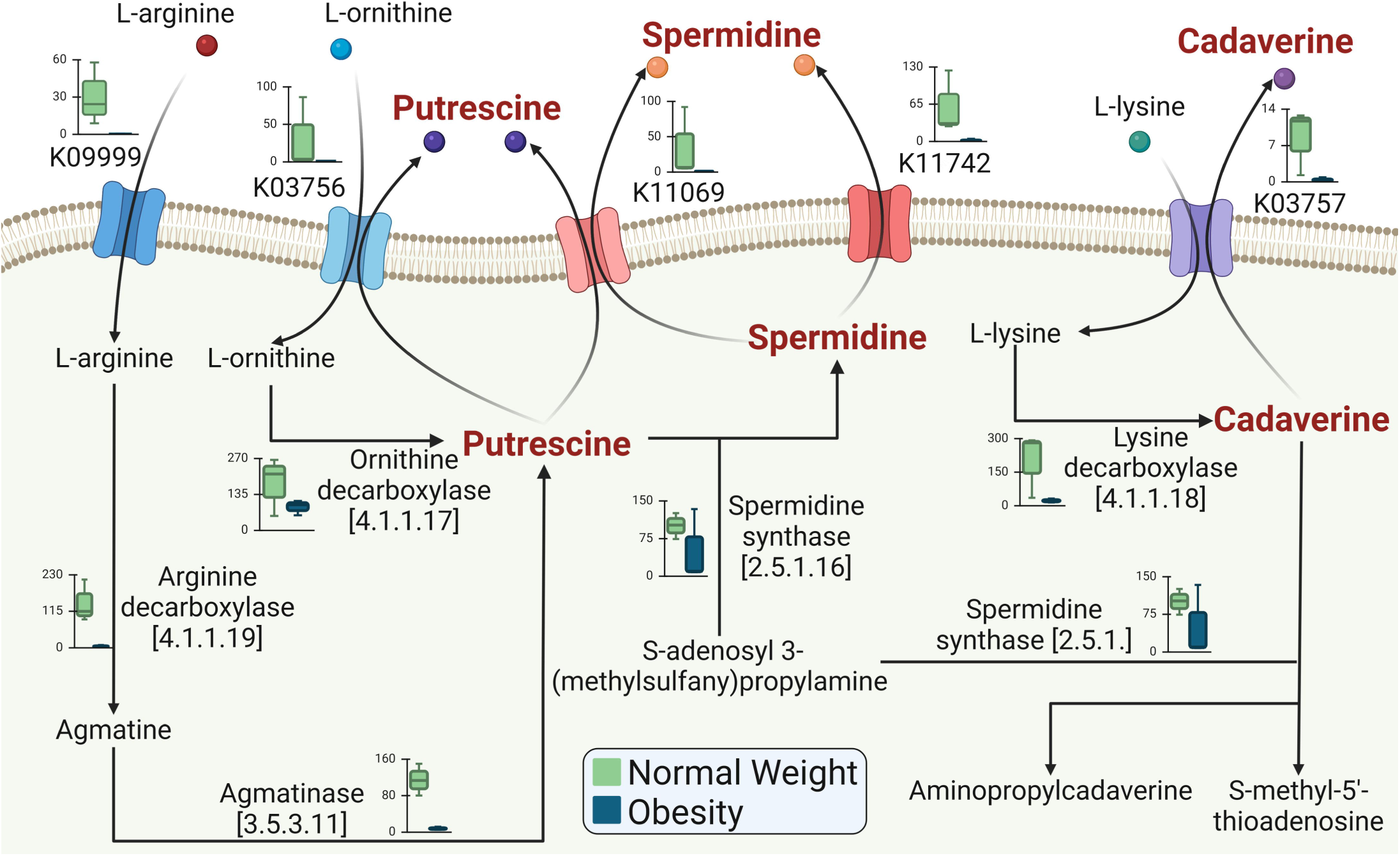
Metabolic polyamine pathway of neonatal microbiome at 48 hr postpartum. Gene abundance comparisons of enzymes and transporters between neonatal microbiomes from normal weight and obese mothers.

In summary, the metagenome of normal weight neonates at M0 is predominantly characterized by pathways related to polyamine and nucleotide biosynthesis, enterobacterial antigen production, and carbohydrate degradation. This metabolism is related to the high predominance of enterobacteria, especially Enterobacter cloacae, whose abundance is decreased in neonates from mothers with obesity. In contrast, the metagenome of this last group is more focused on the degradation of aromatic compounds and energy production through amino acid and ketone biosynthesis pathways.

### Peptidoglycan Biosynthesis and Sulfur-Containing Amino Acid Pathways Are Enriched in the Infant Metagenome of Mothers with Obesity at One Month Postpartum

Similar to taxonomic composition, gut microbial functional profiles in infants diverged only modestly between maternal BMI groups at one month of age compared to the first 48 hours of life. Eighteen pathways exhibited differential abundance (ANCOM-BC, q < 0.05), with 15 pathways enriched in the obesity-associated microbiome and 3 enriched in the normal-weight group (Table S7). Infants born to normal-weight mothers showed elevated activity in pathways linked to pyrimidine biosynthesis and stearate biosynthesis I. In contrast, infants of mothers with obesity harbored microbiome communities enriched for pathways associated with deoxyribonucleotide, stearate II, cysteine and peptidoglycan biosynthesis. Also, for purine nucleobase salvage I, homocysteine-cysteine interconversion, and L-histidine degradation I (Figure 7A).

**Figure 7.**
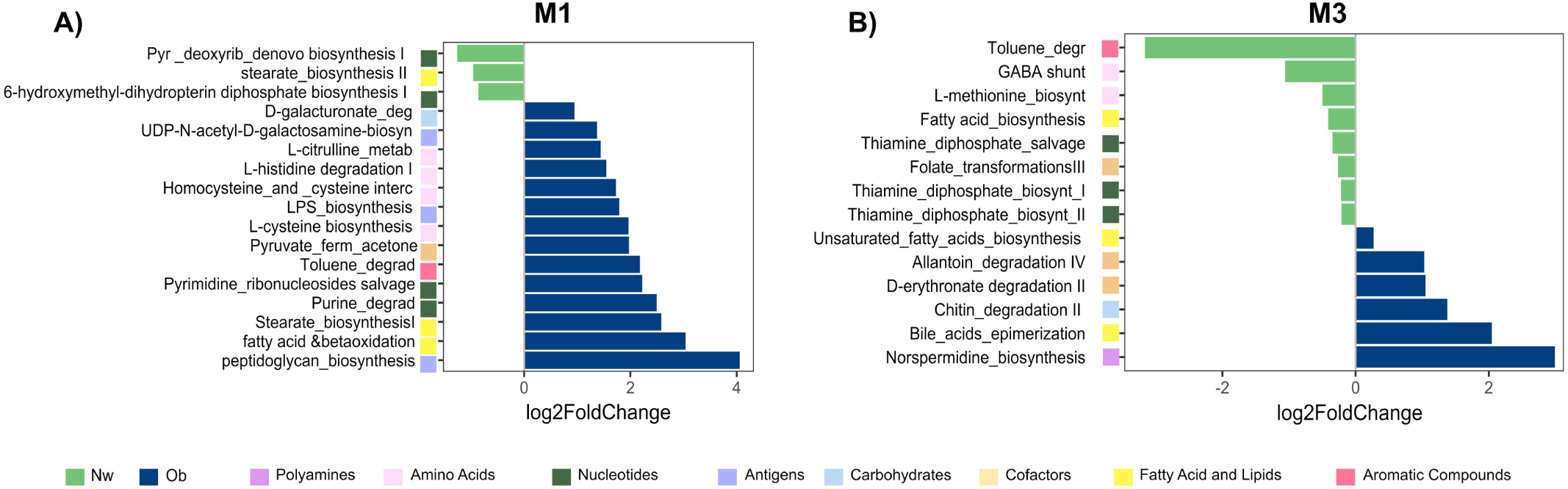
Differential metabolic pathways in the infant microbiome between infants born to normal-weight and obese mothers at (A) 1 month and (B) 3 months postpartum. Horizontal blue bars represent the log2 fold change for pathways enriched in the obesity group, while green bars denote pathways enriched in the normal-weight group, as determined by ANCOM-BC analysis. Pathway categories are indicated by colored rectangles following each pathway name

We observed distinct correlations between maternal oligosaccharide supply and bacterial pathways. Neutral HMOs like LNFPI and LNT, along with 3’-SL, were negatively associated with sulfur-containing amino acid pathways in *Streptococcus mitis* and *S. pneumoniae*. Conversely, 3’-SL and 3’-FL positively correlated with pathways enriched in the obesity group, including those involved in pyrimidine deoxyribonucleotide biosynthesis, stearate production, lipopolysaccharides, and pyruvate-to-acetone metabolism (Figure S3).

### Bile Acid Metabolic Pathway is Elevated in the Metagenomes of Infants Born to Mothers with Obesity at Three Months Postpartum

By the third month postpartum, 16 metabolic pathways exhibited differential gene abundance based on metagenomic sequencing between groups (Table S8). Nine pathways, including thiamine diphosphate salvage, fatty acid biosynthesis, folate transformations, and the GABA shunt (GLUDEG-I-PWY), were overrepresented in the gut microbiomes of normal-weight infants. Conversely, seven pathways such as unsaturated fatty acid biosynthesis (PWY-6284), bile acid epimerization (PWY-6518), and nor spermidine biosynthesis (PWY-6562) were significantly enriched in infants born to mothers with obesity (Figure 7B).

Infants born to mothers with obesity demonstrated elevated microbial metabolic potential for producing secondary bile acids and unsaturated fatty acids. In contrast, the gut microbiomes of normal-weight infants showed stronger associations with pathways linked to folate transformation and GABA shunt activity.

## Discussion

The colonization process in early life is a critical factor with profound immune and metabolic consequences during development. However, our understanding of how maternal obesity influences this process throughout the first months of life remains limited, as most studies focus on single time points in infant development, overlooking the dynamic nature of early microbial colonization. In this study, our findings suggest that infants born to mothers with obesity exhibit an early acquisition of intermediate and late colonizers in the gut microbiome during the first three months of life. Among the factors driving these changes, alterations in human milk components, such as HMOs, appear to play a central role in shaping microbial composition. By examining the associations between HMOs and the infant gut microbiome at specific time points we found that maternal obesity affects both HMO concentrations and their relationships with intermediate and late colonizers in the infant gut. These results suggest that maternal obesity alters early gut colonization patterns and disrupts the natural progression of microbial succession in infancy.

The early neonatal period represents a critical window during which initial bacterial colonization and the dynamic succession of gut microbiota play an essential role in the development of the gut ecosystem and the programming of the immune system (Bokulich et al. 2016; Wang et al. 2020; Wang et al. 2024). During the first 48 hours postpartum, the neonatal gut in most healthy infants is typically dominated by early colonizers such as Proteobacteria, particularly Enterobacteriaceae (Corona-Cervantes et al. 2020; Murphy et al. 2017; Wang et al. 2024; Milani et al. 2017). However, in newborns born to mothers with obesity, we observed a reduced abundance of Proteobacteria in the early postnatal period. This finding aligns with trends reported in previous studies at various time points during the first two weeks after birth (Soderborg et al. 2018; Lemas et al. 2016; Mueller et al. 2016).

In our study, we observed a reduced abundance of metabolic pathways associated with the biosynthesis of immunomodulatory antigens, such as enterobacterial common antigen (ECA) and lipid IVA, in neonates born to mothers with obesity. ECA, a conserved glycolipid in Enterobacteriaceae, triggers cross-reactive antibody production, providing protection against pathogenic strains (Maciejewska et al. 2020; Rai and Mitchell 2020). Similarly, lipid A, the amphipathic glycolipid component of lipopolysaccharides (LPS), produced by strains such as *Klebsiella, Enterobacter* and *Escherichia*, plays a critical role in stimulating immune responses through TLR4/MD-2 signaling pathways (Rai and Mitchell 2020) . TLR4 signaling in epithelial and myeloid cells is essential for inducing immune tolerance particularly during the neonatal period, a time of low microbial diversity and rapid microbial shifts (Ignacio et al. 2024) . Early exposure to lipid A promotes regulatory T cell development and immune tolerance, potentially protecting against immunological disorders later in life (Saha et al. 2022). For example, exposure to LPS from *E. coli* has been shown to protect against the development of type 1 diabetes, but not the *Bacteroides dorei* LPS (Vatanen et al. 2016).

On the other hand, our findings display higher abundances of intermediate colonizers like *Bacteroides*, *Bifidobacterium*, and SCFA-producing taxa such as *Akkermansia*, in newborns from mothers with obesity. This disbalance in the microbiome could have metabolic implications, as the decreased *Gammaproteobacteria*, and the increase in *Clostridia*, *Bacteroides*, and SCFA producers has been linked to metabolic risk in infants of obese mothers (Mueller et al. 2016; Soderborg et al. 2018). While these late bacteria are typically associated with beneficial roles in gut health, such as HMO degradation (Milani et al. 2017; Kostopoulos et al. 2020; Zhang et al. 2021), their premature colonization during the first 48 hours postpartum may disturb the optimal microbial succession pattern. This period is crucial for immune imprinting, and the reduced presence of key early colonizers, such as Proteobacteria could impair myeloid cell development and compromise innate immune function (Soderborg et al. 2018) . This highlights the importance of early exposure to Gram-negative bacteria in training the innate immune system, fostering immune tolerance, and preventing excessive inflammation (Chassin et al. 2010; Ignacio et al. 2024) .

Consistent with our findings at the initial time point, we observed similar shifts in microbial composition that continued to evolve at later sampling points. Our findings suggest that maternal obesity alters this developmental timeline, as infants born to mothers with obesity exhibited a significant enrichment of several *Lachnospiraceae* species as early as one month postpartum. Although *Lachnospiraceae* is generally associated with gut health, early blooms may contribute to obesogenic pathways via SCFA production, potentially influencing energy metabolism (Meehan and Beiko 2014)This aligns with previous studies reporting elevated *Lachnospiraceae* levels in infants of overweight mothers by three to four months postpartum and in neonates of mothers with diabetes (Tun et al. 2018; Forbes et al. 2018; Hu et al. 2013)Additionally, members of this family, such as *Enterocloster* species, have been linked to obesity and type 2 diabetes, suggesting a potential intergenerational microbial signature of obesity (Humphrey et al. 2024). Furthermore, elevated *Lachnospiraceae* abundance during infancy seems to be linked with a higher risk of obesity in later developmental stages such as childhood (Tun et al. 2018) .

Beyond metabolic implications, early expansion of *Lachnospiraceae* species (as *Enterocloster* and *Mediteneibacter*) may also influence immune development, particularly in increasing the risk of allergic conditions such as asthma and atopic dermatitis (Chua et al. 2018; Pirker and Vogl 2024; Korpela et al. 2024) . Early acquisition of *Ruminococcus gnavus* within the first three months may disrupt immune development, as its effects are timing dependent. Breastfeeding has been shown to delay *R. gnavus* expansion until the introduction of non-human-milk foods, allowing for an optimal increase in SCFA production at the appropriate stage (Shenhav et al. 2024). These SCFA-producing microbes help regulate Th2 responses, and their early dysregulation may contribute to immune-related disorders, including asthma (Shenhav et al. 2024; Chua et al. 2018; Chen et al. 2023).

Functional pathway analysis at three months postpartum revealed enhanced bile acid epimerization in infants of mothers with obesity. This process generates iso-bile acids, which are produced by microbes such as *Ruminococcus gnavus* (Sagheddu et al. 2016; Ouyang et al. 2023). Secondary bile acids are potent drivers of early intestinal microbiota maturation, promoting adult-like microbial communities during the critical postnatal period (van Best et al. 2020). With age, interactions between the microbiome and secondary bile acid metabolism intensify, further shaping microbial succession (Tanaka et al. 2020). These findings suggest that maternal obesity may disrupt normative bile acid metabolism, accelerating microbiome maturation through bile acid-modifying taxa. Such early exposure to bile acids or their derivatives could hasten the transition from a neonatal microbiome (dominated by *Bifidobacterium*) to a more diverse, adult-like community.

Another key finding of our study is the consistent negative correlation between neutral non-fucosylated and sialylated HMOs and specific taxonomic groups with maternal obesity at every sampling point. Most of these taxa have been reported to utilize HMOs, either directly or through cross-feeding mechanisms (Sprenger et al. 2022; Kiely et al. 2023; Díaz and Garrido 2024; Masi and Stewart 2022). For example, at the first sampling point, *B. pseudocatenulatum* and *B. longum* were correlated with neutral and sialylated HMOs. These species are well-established primary HMO degraders, employing glycosidases, sialidases, and fucosidases to metabolize HMOs in the infant gut (Kumar et al. 2020; Zhu et al. 2023; Sprenger et al. 2022).

However, most bacteria correlated with HMOs in our results were secondary utilizers, relying on metabolites generated by primary degraders. For instance, *B. breve* ferments fucose, supporting the colonization of *L. reuteri*, while *B. longum* degrades LNT, making HMO-derived metabolites available for *B. pseudocatenulatum* (Xiao et al. 2024) *Bifidobacterium* species also engage in cross-feeding interactions with other gut microbes, such as *L. plantarum*, *Enterococcus faecalis*, and late colonizers like *Ruminococcus gnavus*, *Eubacterium hallii*, and *Akkermansia* (Xiao et al. 2024; Lou et al. 2023) . These species metabolize breakdown products including sialic acid, fucose, acetate, and lactate (Thongaram et al. 2017; Lou et al. 2023). For example, L-fucose produced from fucosylated HMO breakdown by *B. breve* and *B. longum* is utilized by *E. halli* for butyrate production. Additionally, L-fucose availability supports the growth of other microbes encoding fucose catabolic pathways, such as *B. thetaiotaomicron* and *A. muciniphila* (Salli et al. 2021; Schwab et al. 2017; Shuoker et al. 2023)Although the involvement of late colonizers is less common at this early developmental stage, these cross-feeding interactions enable specific gut microbes to thrive, shaping the composition and functional potential of the gut microbiota in later stages of life (Hitch et al. 2022; Kiely et al. 2023).

HMOs not only promote the growth of species and mediate metabolic interactions among them but also restrict the overgrowth of facultative anaerobes like *Enterobacteriaceae* (Torow et al. 2023). The elevated concentration of HMOs in colostrum, compared to later lactation stages, may critically regulate early gut colonizers, including the presence of potential pathobionts which are crucial for establishing a foundation for later microbial succession (Gurung et al. 2024). For instance, LNT structures mimic intestinal epithelial receptors, acting as soluble decoys to block bacterial adhesion. Similarly, sialylated oligosaccharides inhibit pathogen attachment by binding to lectin receptors, while fucosylated HMOs reduce infectivity by impeding pathogen adhesion (Bode and Jantscher-Krenn 2012; Walsh et al. 2020; Gormley et al. 2024; Zhang et al. 2021). Collectively, HMOs shape microbial communities by modulating colonization dynamics and maintaining gut homeostasis.

In addition to HMO modulation, we observed that the metagenomes of neonates born to mothers with normal weight were enriched in genes and pathways associated with polyamine biosynthesis, such as putrescine, spermidine, and cadaverine. These polyamines, primarily derived from arginine and ornithine degradation, are in high demand during neonatal and infant stages due to rapid cell growth (C. Muñoz-Esparza et al. 2024) . We hypothesize that enterobacteria, including *E. coli, Klebsiella aerogenes*, and *Acinetobacter baumannii*, are key producers of polyamines, as they convert arginine to agmatine and generate putrescine via decarboxylation (Wang et al. 2024; Tofalo et al. 2019; Lillie et al. 2024) . Polyamines play a vital role in neonatal gut development by promoting gastrointestinal mucosa growth, reducing mucosal permeability to antigenic proteins, and potentially preventing food allergies (Almeida et al. 2021; Sabater-Molina et al. 2009) . Additionally, they suppress intestinal immunoallergic responses and modulate macrophage polarization. For example, putrescine downregulates M1 gene expression, while spermidine and spermine regulate pro-inflammatory mediator translation in activated macrophages (Atiya Ali et al. 2011; Latour et al. 2020; Ali et al. 2013). The enrichment of polyamine biosynthesis pathways in neonates of normal-weight mothers suggests that early microbial metabolites may directly support neonatal gut development and immune tolerance.

Some limitations must be acknowledged, as the sample size was limited by participant dropout and restricted fecal sample availability, and longitudinal consistency was impacted by incomplete sampling across all time points. While sufficient for an exploratory pilot study, future work should expand cohort sizes and ensure temporal continuity to strengthen generalizability. Additionally, while metagenomic analyses identified functional pathways linked to microbial metabolism, mechanistic validation is required to confirm microbial contributions to host physiology.

Our findings highlight the influence of maternal factors, particularly obesity, on breast milk composition. Reduced HMO levels (2’-FL, 3’-SL, 6’-SL, LNnT and LNFPI) in mothers with obesity may disrupt early postnatal microbiome assembly, with potential consequences for microbial succession and immune system development, including immunological imprinting. These observations underscore the need to prioritize studies of the first days of life, a period critical for gut colonization but poorly explored in current literature. To advance this field, future studies could explore strain-level dynamics to evaluate their role in restoring dysbiosis-associated developmental deficits. Also investigate immune signaling pathways activated by specific HMOs and microbial metabolites. Such efforts could clarify causal relationships between maternal metabolic health, early microbial colonization, and neonatal immunity.

## Acknowledgements

This work was supported by the Challenge-Based Research Funding 2022 of Tecnológico de Monterrey (I008 - IOR001 - C6-T1 – E) to JAGU, CCH and CLC. We thank the National Council for Humanities Science and Technology (CONAHCYT) for providing scholarship to JSGV, Frontera de la Ciencia grant No. CF/2023/G/990 to MB, BIOCODEX Microbiota Foundation to CLC. We would like to thank Juan Pablo Dávila and Aidee Sánchez (from the Center for Protein Development –CIDPRO-) for the support on sample collection for months 1 and 3, to Dr. Ana Lucía Araujo (Resident of Pediatrics) for her support in obtaining the Institutional Review Board approval and thank to the contribution of the Hospital Regional Materno Infantil de Alta Especialidad (HRMIAE) Human Milk Bank personnel for the collection of colostrum samples for the study.

## Author contribution

KCC: Data curation, Formal analysis, Investigation, Supervision, Writing original draft. VHUB: Data curation, Formal analysis, Investigation, Supervision, Writing original draft. JSGV: Data curation, Formal analysis, Methodology, Writing – review & editing.

BJL: Data analysis, Writing – review & editing. NARG: Investigation, Writing-review & editing KarlaCC: Data curation.

FEC: Investigation, Methodology, Writing-review & editing UASV: Investigation, Methodology, Writing-review & editing PARP: Investigation, Writing – review & editing.

JAGU: Supervision, Writing-review & editing.

MB: Data analysis, Writing – review & editing.

CCH: Conceptualization, Funding acquisition, Project administration, Supervision, Writing – review & editing.

CLC: Conceptualization, Data curation, Formal analysis, Funding acquisition, Project administration, Supervision, Writing – original draft, Writing – review & editing.

## Data availability

The raw sequencing data (FASTQ-format reads) generated and analyzed during this study are publicly available in the NCBI Sequence Read Archive (SRA) under the accession number SUB15262662 (BioProject PRJNA1260751).

## Methodology

### Ethics approval for sample collection

This is a longitudinal study conducted in accordance with the ethical principles outlined in the Declaration of Helsinki. Institutional review board approval (approval number: DEISC-PR-190122074; Hospital Regional Materno Infantil de Alta Especialidad de Nuevo León) was obtained. Every participant was provided with details regarding the study, and written consent was acquired from each of them, ensuring the confidentiality of the personal information.

### Experimental model and study participants details

Healthy breastfeeding mothers who had given birth to full-term infants in the Hospital Regional Materno Infantil were recruited between February 2022 and December 2023. The inclusion criteria for participants included women with a full-term pregnancy (≥ 38 weeks); Mexican nationality; healthy; 18 to 49 years of age; that the weight and height record is in the pregnancy control record from the first trimester; patient without a history of diabetes. The exclusion criteria included a prolonged antibiotic exposure exceeding 3 weeks at any stage of pregnancy, antibiotic requirement for more than 24 hours post-delivery, immunosuppressive, or immunomodulatory corticosteroid therapy, and history of feeding disorders or bariatric surgery.

Participants were divided into two groups according to the World Health Organization (WHO) classification of body mass index (BMI): women with obesity (“WO”; BMI ≥ 30 kg/m^2^; n=28) and mothers with normal weight (“NW”; BMI ≤ 25 kg/m^2^; n=20).

Addition, infant weight, and length were recorded during the first, and 3 months and z-scores were obtained according to the WHO databases. Infants were grouped as normal weight (NW) and obesity (OB) according to the pre gestational BMI of their mother.

#### Infant anthropometric measures

Simultaneous weight and length (WL) measurements were collected during periodical pediatric visits by clinicians during the first months of life at birth, 1 and 3 months. The age and sex specific BMI z-score (BMIz) was calculated using. The changes in the BMIz and WLz were calculated by taking the difference in the z-score between time points as described elsewhere.

#### Sample collection and processing

Colostrum samples were collected within the first 48 hours post-partum from a total of 48 mothers by a trained medical team, utilizing sterile gloves for the process. Following a gentle cleansing of the areola with sterile water, when possible, 3 mL of colostrum was obtained by manual expression into a sterile 15 mL polypropylene tube. To ensure the purity of the sample, the initial drops were discarded. After collection, the samples were promptly transported to the laboratory and stored at −20 °C until subsequent processing. The meconium samples from the infants (the first 24 hours) were collected directly from the diaper area with the aid of a sterile tongue depressor and introduced into a 50 mL tube and stored directly at - 20 °C until use.

Home appointments were scheduled for each of the participating mothers and their infants; visits were made at the first and third month postpartum. Human milk samples from the mothers and stool samples from the infants (the first stool of the day) were collected as mentioned above. The samples were immediately placed in a portable cold container and transported to the laboratory in less than 60 minutes and then stored at −20 °C until use.

### HMO quantification

The milk samples were prepared following the steps previously reported by Urrutia-Baca et al., who developed and validated an analytical method for the quantification of HMOs (Urrutia-Baca et al. 2024) . An Acquity UPLC-MS/MS (Waters, Milford, MA) equipped with Micromass Quattro Premier XE Mass Spectrometer (Waters, Milford, MA) was used to analyze the HBM samples with the following specifications: run time, 14 min; Aquity UPLC column (2.1 x 100 mm, 1.7 µm, 40°C); 0.400 mL/min flow; argon (collision gas) and nitrogen (running gas); sample manager (10°C, 2 µL injection volume per sample); 10 mM ammonium formate in LC-MS water and acetonitrile (100%) and as mobile phase A and B, respectively. The following linear UPLC gradient was used: 02 min from 98% to 95% B, 23 min to 65% B, 38.5 min to 55% A, 8.511 min to 55% A, 11-11.2 min to 98% B, 11.2-14 min to 98% B. In addition, the mass spectrometer was operated in negative-ion, multiple reaction monitoring (MRM) mode.

### Metagenomic DNA extraction and sequencing

To investigate whether the gut microbiome of infants is influenced by maternal HMO supply, a pilot study was conducted using fecal samples from a subset of 26 infants. These infants were sampled at birth, and at 1 and 3 months of age. Fecal samples were collected immediately after defecation from the diaper into sterile collection tubes. The samples were then aliquoted into sterile tubes under a biological safety cabinet and stored at −20°C until further processing.

Genomic DNA from bacterial cells was extracted from 200 mg of stool samples using the Favorgen FavorPrep Stool DNA Isolation Kit (catalog number: FASTI 001-1) according to the manufacturer’s instructions. DNA concentrations were measured using the Qubit Flex Fluorometer from Thermo Fisher Scientific. The extracted DNA was then used to prepare metagenomic shotgun sequencing libraries with the Illumina DNA Prep kit and custom 10 bp unique dual indices (UDI) from Integrated DNA Technologies (IDT), aiming for an average insert size of 280 bp. Sequencing was carried out on an Illumina NovaSeq X Plus platform using 2×151 bp paired-end reads in one or more multiplexed, shared-flow-cell runs, following standard Illumina protocols. Demultiplexing, quality control, and adapter trimming were performed using bcl-convert1 (version 4.2.4). A reagent blank control, using an extraction without a sample, was included as a negative control to check for contamination.

### Metagenomic Data Quality Control Quality and Filtering

Quality trimming of the reads and removal of residual adapters were performed using the Trimmomatic v0.39 software with the following parameters: -phred33, MINLEN:100, SLIDINGWINDOW:4:20, and MINLEN:35. Sequencing reads were identified and filtered in each sample mapping to the human genome reference assembly HG38 GRCh38.p12 using BOWTIE2 with default settings and filtered using SAMtools. Following data quality control and removal of host sequences, taxonomic profiling was conducted using Kraken2 with default parameters. Metabolic pathway analysis and gene abundance quantification were performed using HUMAnN3, with results reported in reads per kilobase (RPKs) and subsequently normalized to copies per million (CPM). The UniRef90 database annotations and MetaCyc Reactions were used as reference databases. Specific genes that were more abundant or less abundant in either group, or that mapped to specific COGs having statistically significant differences in representation between stages.

### Statistical Analysis

Microbiome bacterial abundance and alpha diversity metrics (including Observed, ACE, Chao1, Shannon, Simpson, and Fisher indices) were analyzed using the Phyloseq package (v2.6.4). Stacked bar plots and heat maps were created using ggplot2 (v3.5.1). Beta diversity was calculated using the Bray-Curtis dissimilarity index and analyzed through permutation analysis of variance (PERMANOVA). Differential abundance analysis of species, functional pathways, and genes was performed using the q2-composition ANCOMBC method in QIIME 2024, with default settings and significance thresholds set at p<0.05. Multiple comparisons were adjusted using the Holm method, considering q<0.05.

Linear mixed-effects models (LMMs) were employed to assess the association between HMO abundance and various clinical, biochemical, and sociodemographic factors of the mother and infant (including maternal age, gestational weeks, delivery mode, and neonatal sex) for longitudinal data. Each fixed and explanatory variable was fitted using the lme4 package in R (Bates et al. 2015). These covariates were selected based on biological plausibility and prior evidence of their impact on gut microbiome composition in large birth cohorts. All models included the stage or month of sampling as a random effect, and interactions with the sampling point were also analyzed. A p-value of <0.05 was considered statistically significant.

Correlations between functional pathways, taxonomic composition, HMO concentrations, maternal parameters, and infant developmental parameters were analyzed using Spearman correlation and visualized as heatmaps with the microbiome package. Only taxa with p-values of <0.1 and <0.05 were retained for visualization in the heatmaps.

