## Supplementary Figures for "Maternal obesity alters human milk oligosaccharides content and correlates with early acquisition of late colonizers in the neonatal gut microbiome"

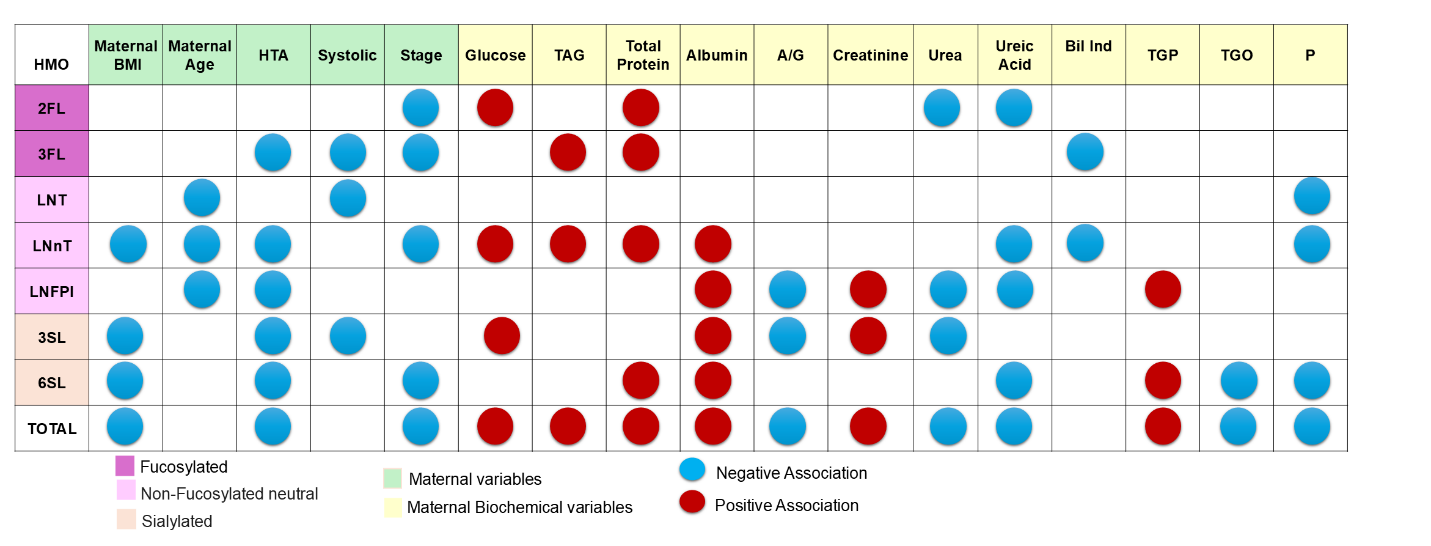

**Figure S1.** *Association of Maternal Variables with HMO Concentrations in Human Milk.* Linear mixed-effects models were applied to data from mothers who provided human milk samples. Abbreviations: HTA = Hypertension, Systolic = Systolic Pressure, Occupation = Domestic worker, IB = Indirect Bilirubin, TAG = Triglycerides, FA = Fatty Acids, TGO = Glutamic Oxaloacetic Transaminase, TGP = Glutamic Pyruvate Transaminase, A/G = Albumin-Globulin Index. The graphical representation is based on the beta coefficients obtained using the LMM in the *lme4* package. Full numeric data are available in Supplementary Tables 2 and 3. For clinical and sociodemographic variables, data from all three lactation stages were used, while for biochemical variables, only M1 and M3 samples were considered.

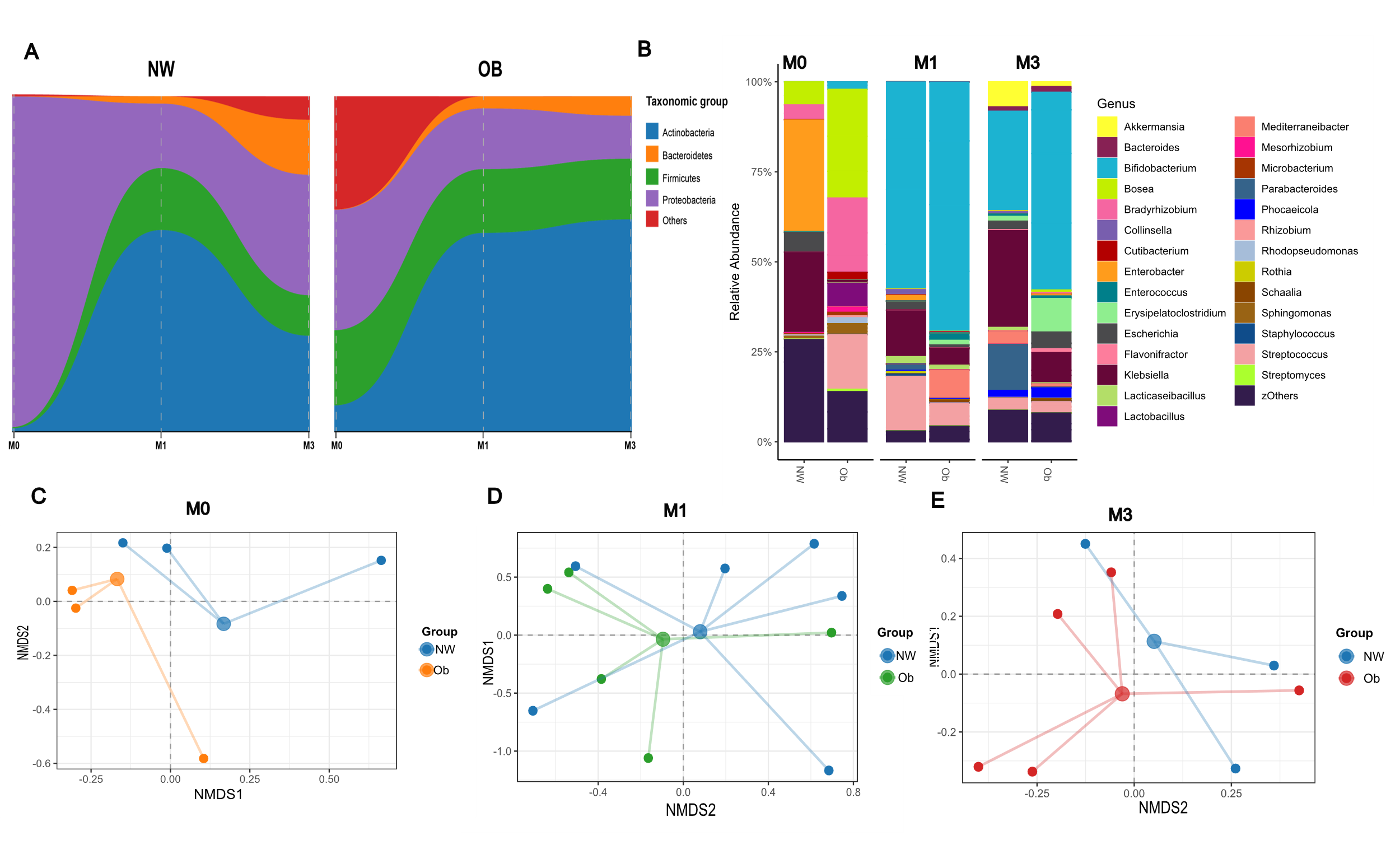

**Figure S2.** *Taxonomic Description of Infant Fecal Microbiota Composition and Abundance During the First 3 Months of Life According to Maternal BMI.* (A) Relative abundance of bacteria at the phylum level and (B) genus level at birth (M0), one month (M1), and three months (M3) postpartum. Taxa are ordered in the legend to correspond with the color sequence in the figure, representing bacteria with >1% average relative abundance in the infants' feces. Samples are clustered based on maternal BMI. Adonis pairwise comparisons: NW_OB (p=0.200, R²=0.258) at M0, NW_OB (p=0.754, R²=0.061) at M1, and NW_OB (p=0.393, R²=0.147) at M3.

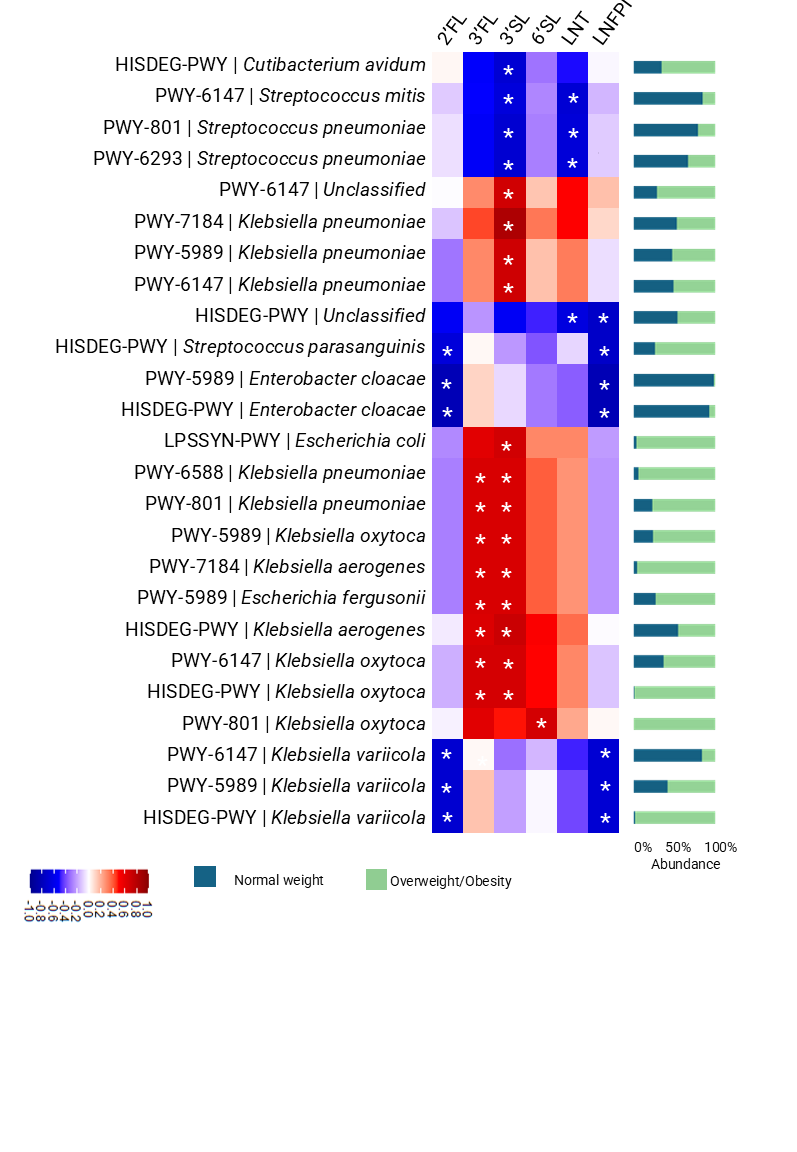

**Figure S3.** *Correlation functional metabolic pathways at 1 month postpartum in infant microbiome with Human milk oligosaccharides concentration*. The Spearman correlation index was employed to establish correlation. Blue shows negative correlation and red for positive. Stars represent a p value <0.05. Stacked Bar plots represent the relative abundance of the stratified pathway according to the maternal BMI.

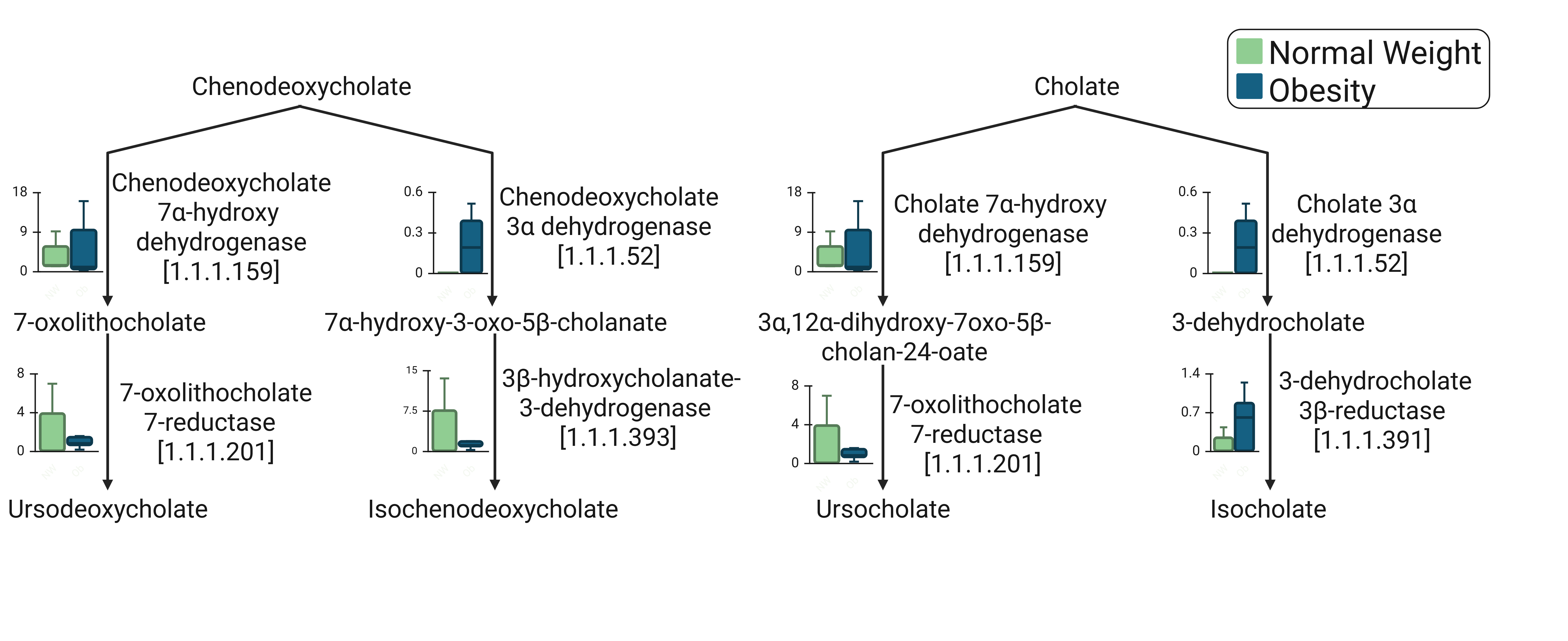
**Figure S4.** *Epimerization of biliary acids pathway in infants at 3 months postpartum.*
